# A framework for defining mesenchymal cell types associated with murine periosteal and endosteal bone

**DOI:** 10.1101/2023.11.17.567528

**Authors:** Intawat Nookaew, Jinhu Xiong, Melda Onal, Cecile Bustamante-Gomez, Visanu Wanchai, Qiang Fu, Ha-Neui Kim, Maria Almeida, Charles A. O’Brien

## Abstract

Single-cell RNA sequencing has led to numerous novel designations for mesenchymal cell types associated with bone. Consequently, there are now multiple designations for what appear to be the same cell type. In addition, existing datasets contain relatively small numbers of mature osteoblasts and osteocytes and there has been no comparison of periosteal bone cells to those at the endosteum and trabecular bone. The main goals of this study were to increase the amount of single cell RNA sequence data for osteoblasts and osteocytes, to compare cells from the periosteum to those inside bone, and to clarify the major categories of cell types associated with murine bone. To do this, we created an atlas of murine bone-associated cells by harmonizing published datasets with in-house data from cells targeted by Osx1-Cre and Dmp1-Cre driver strains. Cells from periosteal bone were analyzed separately from those isolated from the endosteum and trabecular bone. Over 100,000 mesenchymal cells were mapped to reveal 11 major clusters designated fibro-1, fibro-2, chondrocytes, articular chondrocytes, tenocytes, adipo-CAR, osteo-CAR, pre-osteoblasts, osteoblasts, osteocytes, and osteo-X, the latter defined in part by *Postn* expression. Osteo-X, osteo-CAR, and pre-osteoblasts were closely associated with osteoblasts at the trabecular bone surface. Wnt16 was expressed in multiple cell types from the periosteum but not in any cells from endocortical or cancellous bone. Fibro-2 cells, which express markers of skeletal stem cells, localized to the periosteum but not trabecular bone in adult mice. Suppressing bone remodeling eliminated osteoblasts and altered gene expression in pre-osteoblasts but did not change the abundance or location of osteo-X or osteo-CAR cells. These results provide a framework for identifying bone cell types in murine single cell RNA sequencing datasets and suggest that osteoblast progenitors reside near the surface of remodeling bone.

**Author Summary:** The skeleton of vertebrate animals is produced by bone forming cells known as osteoblasts. New osteoblasts are continually produced throughout life to maintain the skeleton. However, the identity of the cells from which new osteoblasts are derived is not clear. Single cell RNA sequencing provides a gene expression profile of individual cells and allows them to be grouped into clusters of cells with similar profiles. In many cases, distinct clusters represent distinct cell types. Recent single cell RNA sequencing studies of mouse bones by several different laboratories have identified many novel cell types, some of which may be osteoblast progenitors. In many cases, different laboratories have created different names for what appear to be the same cell type. To help clarify this situation, we performed single cell RNA sequencing studies and combined our results with those from several published studies to create a harmonized map of the cell types associated with mouse bones. We also identified gene signatures for each cell type that can be used to improve the consistency of cell type designation in future single cell RNA sequencing studies of mouse bone.

## Introduction

To develop a comprehensive understanding of how different cell types work together to create and maintain the vertebrate skeleton, it is essential to clearly identify the cell types involved. Initially, the cells that produce and maintain bone were defined by their morphology and location in histological sections (1–3), followed by studies using cultured cells, colony-forming assays, and transplant assays (4–8). Analysis of gene expression, either in isolated cells, primary cell cultures, or continuous cell lines, has also been used to define bone cell types (4). Moreover, detection of gene products in histological sections, via antibodies, in situ hybridization, or enzymatic activity, underpins much of our current knowledge of skeletal cell types. Conditional gene inactivation in cell types defined by expression of particular genes has also provided a wealth of information regarding the function of different genes and the cell types that express them (9).

Even with this abundance of knowledge, important questions remain regarding how different skeletal cell types are related to one another and regarding their full range of functions. For example, while it is clear that beta-catenin in a subset of cells targeted by Cre driver strains such as Osx1-Cre is essential for osteoblast specification and differentiation (10–13), the location of these cells in the bone microenvironment and their relationship to other cells of the mesenchymal lineage remain unclear. Similarly, Dmp1-Cre driver strains are often used to study the role of particular genes in osteocytes (14, 15). However, interpretation of results from such studies is complicated by the fact that the full range of cell types targeted by Dmp1-Cre strains is not known (15, 16). These situations highlight the need for more precise definition of the cell types involved in skeletal formation and maintenance and which of these cells are targeted by various Cre driver strains.

Single cell RNA sequencing (scRNA-seq) provides a quantitative analysis of mRNA transcripts in hundreds to thousands of individual cells in a single experiment (17). Based on the transcriptomic profile of individual cells, they can be grouped or clustered so that cells with similar profiles are grouped together. Combining this information with the known specificity of gene expression obtained from other approaches can be used to identify or define clusters of cells in scRNA-seq datasets as a specific cell type. Several studies over the last 5 years have used scRNA-seq to define cells associated with murine bone, which include cells of the mesenchymal lineage, endothelial cells, and hematopoietic cells (18–26). Many of these studies have focused on defining types of mesenchymal stromal cells, which are thought to contain mesenchymal stem cells, as well as the progenitors of chondrocytes, osteoblasts, and adipocytes.

The first goal of this study was to increase the amount of sequencing data from osteoblasts and osteocytes because previous scRNA-seq studies included only small numbers of these cell types. A second goal was to better define the clusters, or cell types, associated with murine long bones. To accomplish these goals, we combined our results, which sequenced cells targeted by Osx1-Cre and Dmp1-Cre transgenes, with several published scRNA-seq datasets to develop a more comprehensive and uniform definition of mesenchymal cell types associated with murine long bones. Our results define 11 broad categories of cell types and identify which of these are targeted by Osx1-Cre and Dmp1-Cre in the periosteal and endosteal compartments of long bones.

## Results

### Isolation of cells using Osx1-Cre and Dmp1-Cre

To increase the abundance of osteoblasts and osteocytes in scRNA-seq datasets, we isolated cells targeted by Osx1-Cre or Dmp1-Cre transgenes by crossing these strains with Ai9 Cre reporter mice (27). We focused on cells associated with endosteal and trabecular bone in adult mice by removing the periosteum, epiphyses, and bone marrow from tibias and femurs followed by sequential incubation of the remaining bone fragments with collagenase and EDTA. Cells expressing high levels of tdTomato were isolated by FACS and subjected to scRNA-seq using the 10X Chromium platform (**Fig. S1**). For brevity we will refer to these cells as endosteal but recognize that this preparation contains cells from cancellous bone as well as the endosteum. From two independent isolations from Dmp1-Cre;Ai9 mice we obtained sequence from a total of 3,845 cells (Fig. S1). From a single isolation from Osx1-Cre;Ai9 mice, we obtained sequence from 6,071 cells. In a separate set of experiments, we also isolated targeted cells from the periosteum and cortical bone surface of long bones and obtained sequence from 4,106 cells targeted by Dmp1-Cre and 15,113 cells targeted by Osx1-Cre (Fig. S1). Initial cluster analysis revealed that hematopoietic and endothelial cells made up only a small proportion of the sequenced cells from each isolation (Fig. S1). The presence of hematopoietic and endothelial cells in these preparations likely represents contamination or rare recombination of the Ai9 locus in these cells, with the latter idea supported by detectable levels of the tdTomato transcript in many cells in these clusters (Fig. S1).

The majority of the cells from each isolation appeared to represent different types of mesenchymal cells based on expression of known stromal cell and osteoblastic marker genes, such as *Col1a1* and *Pdgfra* (Fig. S1). One approach to define cell types in our analysis would be to pool the results from the different experiments and perform a cluster analysis using only our own data. However, we and others have noted that scRNA-seq studies from independent laboratories often identify what appear to be the same or similar cell types (clusters) but use novel designations (23, 28). In order to avoid propagating this growing complexity in cell designations, we sought to place our results into the context of published work by performing a pooled analysis including our data as well as multiple published datasets. To accomplish this, we reprocessed published datasets and performed canonical correlation analysis-based procedures to normalize our data with those from 9 studies that focused on cells associated with murine bone and that used the 10X Genomics platform (18–26). By combing our new datasets with the published work, approximately 190,000 high quality cells were included and projected on UMAP coordinates to uncover the structure of clusters representing different cell types (**Fig. S2**).

Even though our datasets included relatively small numbers of hematopoietic cells, the published datasets we used for harmonization contained sufficient numbers of these cell types so that definitive clusters representing all major hematopoietic cell types were apparent (Fig. S2). Similarly, clusters for endothelial cells, pericytes, and Schwann cells were also readily identified (Fig. S2). Because the goal of our study was to better define mesenchymal cell types, we focused the remainder of the study on this group of cell types.

### Definition of mesenchymal clusters

Harmonization of the 9 published datasets with our data produced a UMAP of 100,000 mesenchymal cells in 11 major clusters (**Fig. 1A-C**). Because of the large number of cells included, and because the cells were isolated using a variety of methods by independent laboratories (**Fig. S3**), this reference map may be useful as a guide for the analysis of future scRNA-seq experiments using murine bone-associated cells.

**Figure 1.**
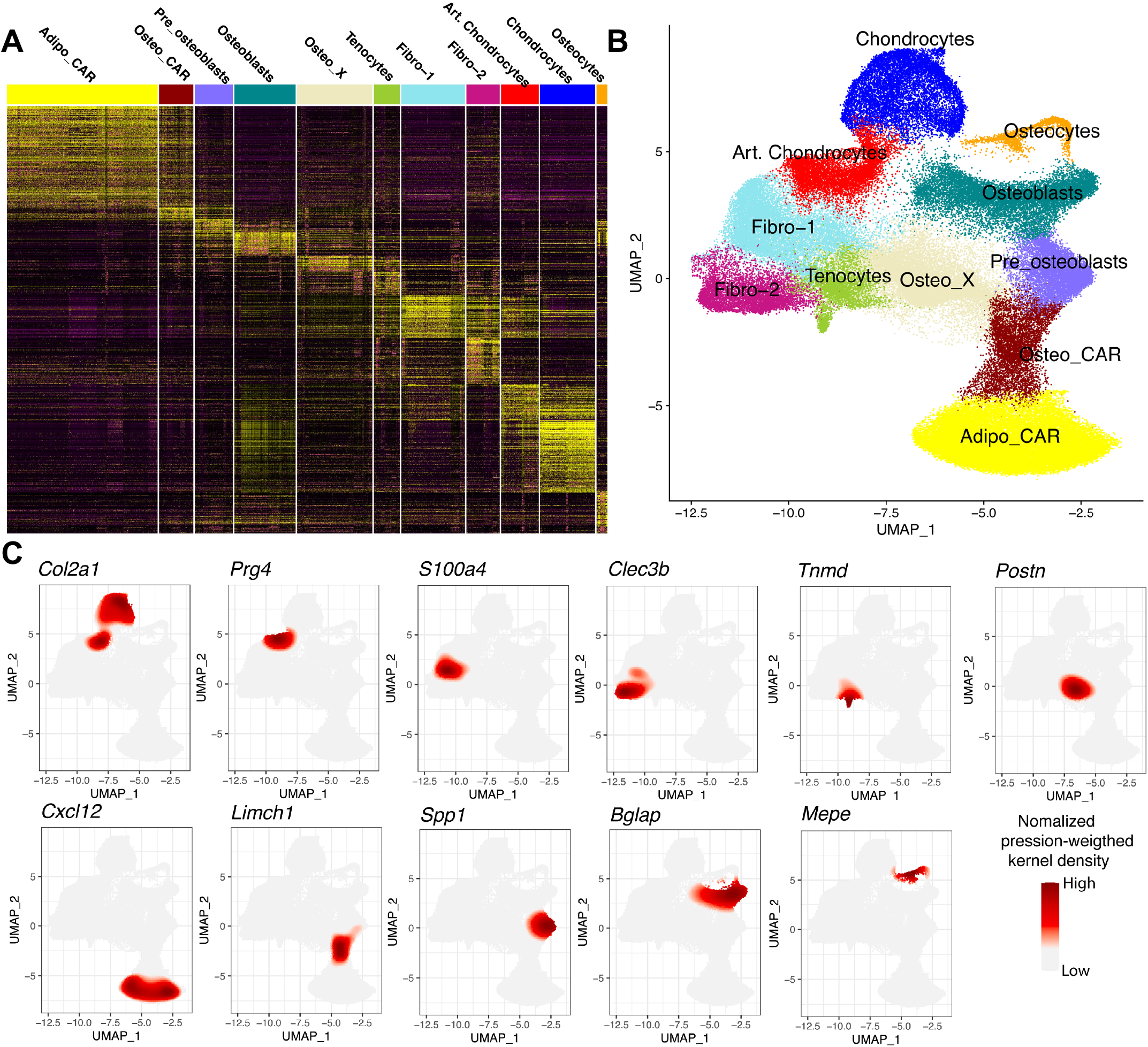
Harmonization of new and existing bone mesenchymal cell datasets. Results shown in this figure were derived from single cell RNA-seq analysis of 100,000 mesenchymal cells, of which 26,535 were isolated and sequenced in the present study. **A.** Heatmap of transcript abundance for the major genes driving definition of 11 mesenchymal cell clusters. Cell names and color codes are shown at the top. **B.** UMAP representation of the 11 major clusters of mesenchymal cells. Names and color codes same as in A. **C.** UMAP feature plots showing expression of representative cell type-specific transcripts. Red = high expression.

The following is a brief description of each cluster. We endeavored to use cluster nomenclature that is consistent with the earliest published study describing that cell type (18, 19). It is important to note that the number of cells in each cluster does not necessarily represent the relative abundance of that cell type in bone. The reason for this is that the procedures used to isolate the cells in each study selected for and against different cell populations (Fig. S3). In addition, some cell types, such as mature adipocytes, may not be represented at all due to their inability to survive the isolation processes that were used.

#### Chondrocytes

Cells in the chondrocyte cluster express high levels of *Col2a1*, *Sox9*, as well as the genes *Snorc*, *Fxyd2 and Matn3* (**Figs. 1, 2 and S4**). This cluster likely contains many stages of the chondrocytes present in long-bone growth plates as evidenced by subpopulations expressing *Col10a1*, *Ihh*, and *Pthlh* (**Fig. S5**).

#### Articular Chondrocytes

Lubricin, encoded by *Prg4*, is highly expressed in articular chondrocytes and synovial fibroblasts (29). Therefore, a cluster expressing high levels of *Prg4*, as well as *Cytl1*, *Cpe*, and *Crispld1* was designated as articular chondrocytes (Figs. 1, 2 and S4). *Fibro-1 -* Cells in this and the next cluster have been designated fibro-1 and fibro-2 based on the high expression of genes associated with bone-associated fibroblasts, such as *S100a4* and *Dcn* (19). We acknowledge that the term fibroblast is vague, but its use here reflects the uncertainty regarding the identity of the cell types in these clusters. In addition to *S100a4*, the fibro-1 cluster is defined by expression of *Angptl7*, *Cilp2*, and *Anxa8* (Fig. 2 and S4).

**Figure 2.**
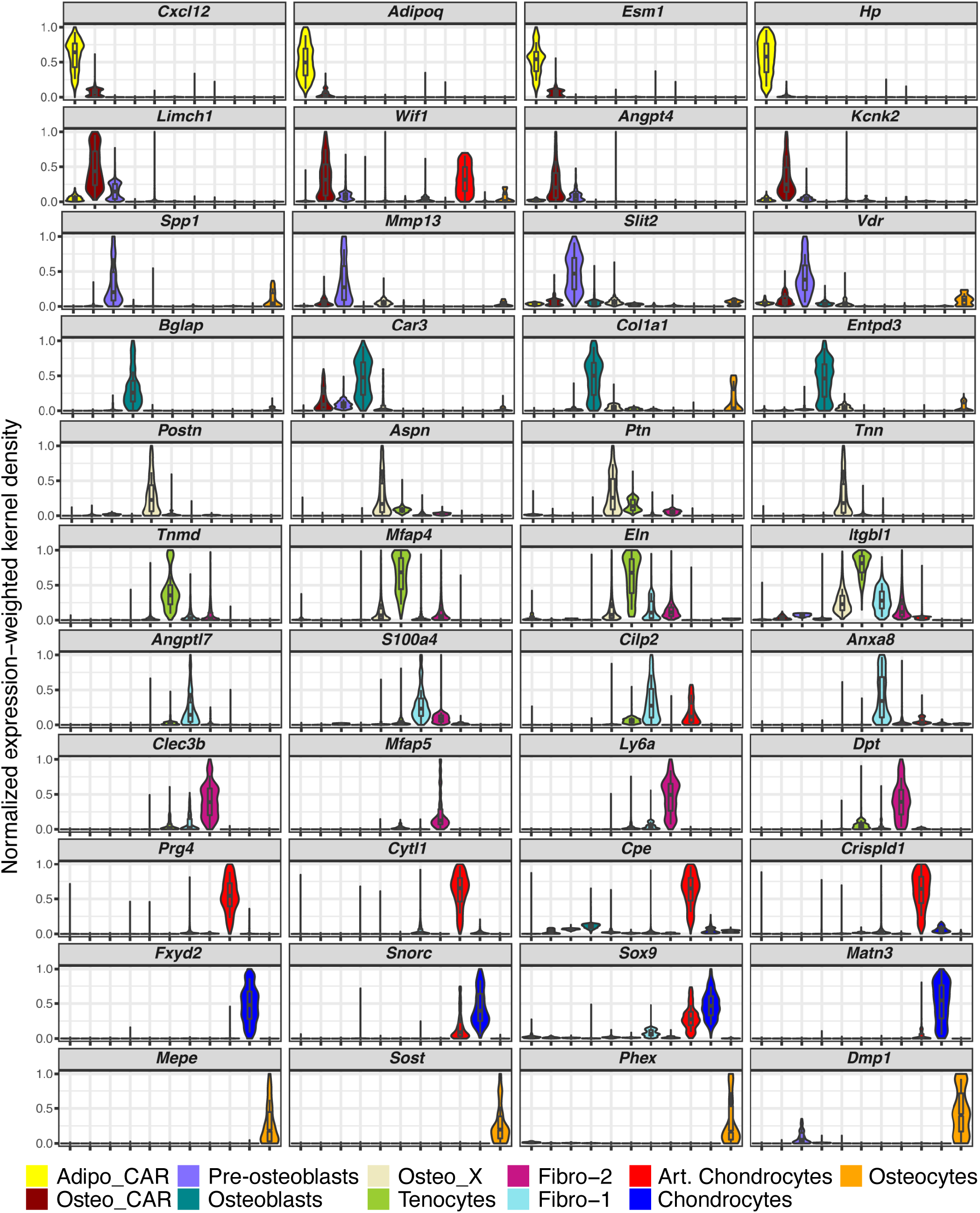
Major transcripts defining mesenchymal cell clusters. Violin plots showing expression levels of four major transcripts defining each of the 11 major mesenchymal cells clusters using the same dataset used in figure 1. The box plots show the maximum value, first quartile value, median value, third quartile value, and the minimum value. Cell names and color codes are the same as in figure 1.

#### Fibro-2

Many of the cells in the fibro-2 cluster express *Ly6a* and *Cd34*, which are often used as markers of progenitor cells (19, 30). As noted by others, mesenchymal lineage cells that express these genes may represent mesenchymal stem cells or other mesenchymal progenitors (19). Cells in this cluster also express *Clec3b* and *Mfap5*, which, unlike *Ly6a* and *Cd34*, are not highly expressed by endothelial or hematopoietic clusters, suggesting that they may be more specific markers for skeletal mesenchymal progenitors (Fig. 2 and S4).

#### Tenocytes

Tenomodulin (*Tnmd*) is one of the most tendon-specific gene products (31) and is a marker that defines a cluster that likely contains tenocytes (Fig. 2 and S4). Other genes enriched in this cluster include *Mfap4*, *Eln*, and *Itgbl1*.

#### Osteo-X

We have used the novel, and likely temporary, designation of Osteo-X to define a cell population that highly expresses *Postn* (Periostin), *Aspn* (Asporin), and *Tnn* (Tenascin-W) (Fig. 2 and S4). These genes are highly expressed in periosteal cells (32–34), suggesting that many of the cells in this cluster originated from the periosteum. Consistent with this idea, many more osteo-X cells were isolated from Osx1-Cre mice when periosteal tissue was included, compared to bones stripped of periosteum (**Fig. 3A**).

**Figure 3.**
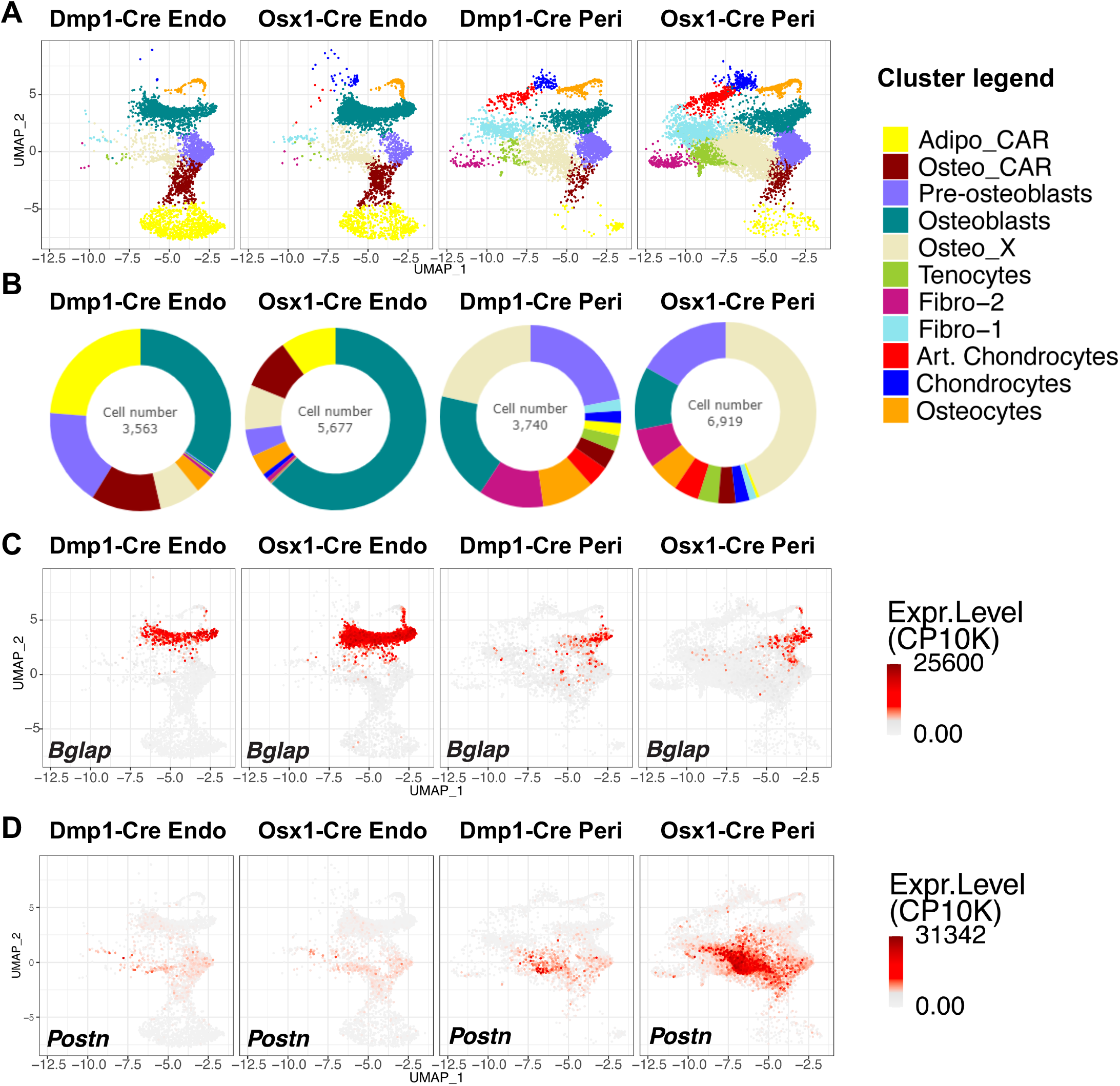
Osx1-Cre targets periosteal osteo-X cells better than Dmp1-Cre. **A.** UMAPs of the cells targeted by Dmp1-Cre or Osx1-Cre in endosteal and trabecular bone (Endo) or in periosteal bone (Peri). Cell names and color code are the same as in Figures 1 and 2. **B.** Pie charts depicting the relative number of cells in each cluster targeted by Dmp1-Cre or Osx1-Cre at each site. **C-D.** UMAP feature plots showing the abundance of *Bglap* and *Postn* transcripts in the same cell preparations shown in panel A.

#### Adipo-CAR cells

High expression of the chemokine Cxcl12 has been used to define a population of stromal cells known as Cxcl12 abundant reticular (CAR) cells (35). Baccin and coworkers classified CAR cells into two distinct subsets based on their gene expression profile, adipo-CAR and osteo-CAR (18). In our analysis, adipo-CAR cells express the adipogenic marker *Adipoq* in addition to high expression of the stromal marker *Cxcl12* (Fig. 2 and S4). However, these cells express relatively low levels of the osteoblastic markers *Col1a1* and *Bglap* (Fig 2 and S4). Adipo-CAR cells also express high levels of *Esm1* and *Hp* (Fig 2 and S4).

#### Osteo-CAR cells

Cells in the lower region of the cluster designated osteo-CAR express stromal markers such as *Cxcl12* and *Adipoq*, whereas cells in the upper region express osteoblastic markers such as *Col1a1* and *Bglap* (Fig. 2 and S4). Transcripts that define this cluster include *Limch1*, *Wif1*, *Angpt4*, and *Kcnk2* (Fig. 2 and S4).

#### Pre-osteoblasts

A cluster that expresses high levels of *Spp1*, also known as osteopontin, and *Mmp13* was designated as pre-osteoblasts due to the continuity with the osteoblast cluster and expression of osteoblast marker genes in the upper portion of this cluster (Fig. 2 and Fig. S4). Other genes enriched in this cluster include *Slit2* and *Vdr*. Additional support for the idea that these cells represent osteoblast progenitors will be presented in later sections of this study.

#### Osteoblasts

The osteoblast cluster was identified by high expression of the genes encoding osteocalcin (*Bglap*) and type 1 collagen (*Col1a1*) but lacking expression of known osteocyte-specific genes such as *Mepe* and *Sost* as well as lower levels of *Dmp1* (Fig. 2 and Fig. S4). Cells in the osteoblast cluster also express high levels of *Car3* and *Entpd3*, as noted by Tikonova and colleagues (20) (Fig. 2 and Fig. S4).

#### Osteocytes

This cluster expresses genes previously shown to be either produced specifically by osteocytes or highly-expressed by osteocytes, including *Mepe*, *Sost*, *Phex*, and *Dmp1* (Figure 2 and Fig. S4) (36). These cells continue to express high levels of gene products characteristic of osteoblasts including *Col1a1* (Fig. 2 and Fig. S4).). We compared the gene expression profile of our osteocyte dataset to previously published scRNA-seq (26, 37) and bulk RNA-seq (38) osteocyte profiles (**Fig. S6**). Only three transcripts were present in all four profiles: *Dkk1*, *Dmp1*, and *Irx5*. Notably, two of the most osteocyte-specific transcripts, *Sost* and *Mepe*, were present in only two of the profiles, ours and the one from Youlten et. al (38).

Of the previously published scRNA-seq studies used for harmonization, only the study by Wang et al. reported a distinct cluster labeled as osteocytes (26). One reason for this may be the technical difficulty associated with isolating these cells from mineralized bone. Nonetheless, it is likely that such cells were present in most of the previous studies but were not uncovered as a separate cluster due to their small representation. Our integrated analysis uncovered 1,840 osteocytes in total, including 1,100 osteocytes from our experiments.

### Osx1-Cre targets more osteo-X cells than Dmp1-Cre

Having defined the clusters, we next compared the harmonized scRNA-seq results of endosteal cell preparations from Osx1-Cre and Dmp1-Cre and found that these driver strains labeled very similar cell populations, including osteo-X, osteo-CAR, adipo-CAR, pre-osteoblasts, osteoblasts, and osteocytes (**Fig. 3A-B**). A small number of fibro-1, fibro-2, and chondrocytes were also labeled, but together these cell types represented less than 5**%** of the cells labeled by either transgene. A striking difference between the two transgenes was the percentage of cells in the osteoblast cluster, which represented approximately 30% of the Dmp1-Cre targeted cells but approximately 60% of the cells targeted by Osx1-Cre (**Fig. 3A-C**). The significance of this observation is unclear but suggests that there may be a population of osteoblasts that are not targeted by Dmp1-Cre.

In the periosteal cell preparations, the greatest difference between Osx1-Cre and Dmp1-Cre was targeting of the osteo-X cluster (Fig. 3A-B). While the percentages of osteo-X, pre-osteoblasts, and osteoblasts were similar to one another in Dmp1-Cre targeted cells, the percentage of osteo-X cells targeted by Osx1-Cre was more than 3-fold greater than either pre-osteoblasts or osteoblasts (Fig. 3B). This preferential targeting was also evident in the expression pattern of the osteo-X marker transcript *Postn* (**Fig. 3D**). These results suggest that Osx1-Cre targets osteo-X cells more effectively than Dmp1-Cre does.

Because the difference in osteo-X targeting was not observed in the endosteal preparations, we explored whether there were significant differences in gene expression that might distinguish osteo-X cells in the periosteum from those at the endosteum. For this comparison, we also included published datasets from calvaria and calvarial sutures (24, 25), both of which contain significant percentages of osteo-X cells (**Fig. 4A**). Comparison of osteo-X cells from the periosteum to osteo-X cells from the other sites revealed that a handful of transcripts, including the one encoding *Wnt16*, were greatly enriched in periosteal cells (**Fig. 4B**). In situ hybridization confirmed expression of *Wnt16* exclusively in periosteal cells (**Fig. 4C-D**). However, expression of *Wnt16* at the periosteum was not confined to osteo-X cells, but also occurred in several other cell clusters (**Fig. 4E**). Thus, *Wnt16* expression does not represent a unique property of periosteal osteo-X cells but rather a property of periosteal cells in general. The small level of differential gene expression between osteo-X cells from the periosteum versus the endosteum suggests that the preferential targeting of these cells by Osx1-Cre at one site but not the other is unrelated to differences in gene expression but instead may result from greater abundance of these cells at the periosteum.

**Figure 4.**
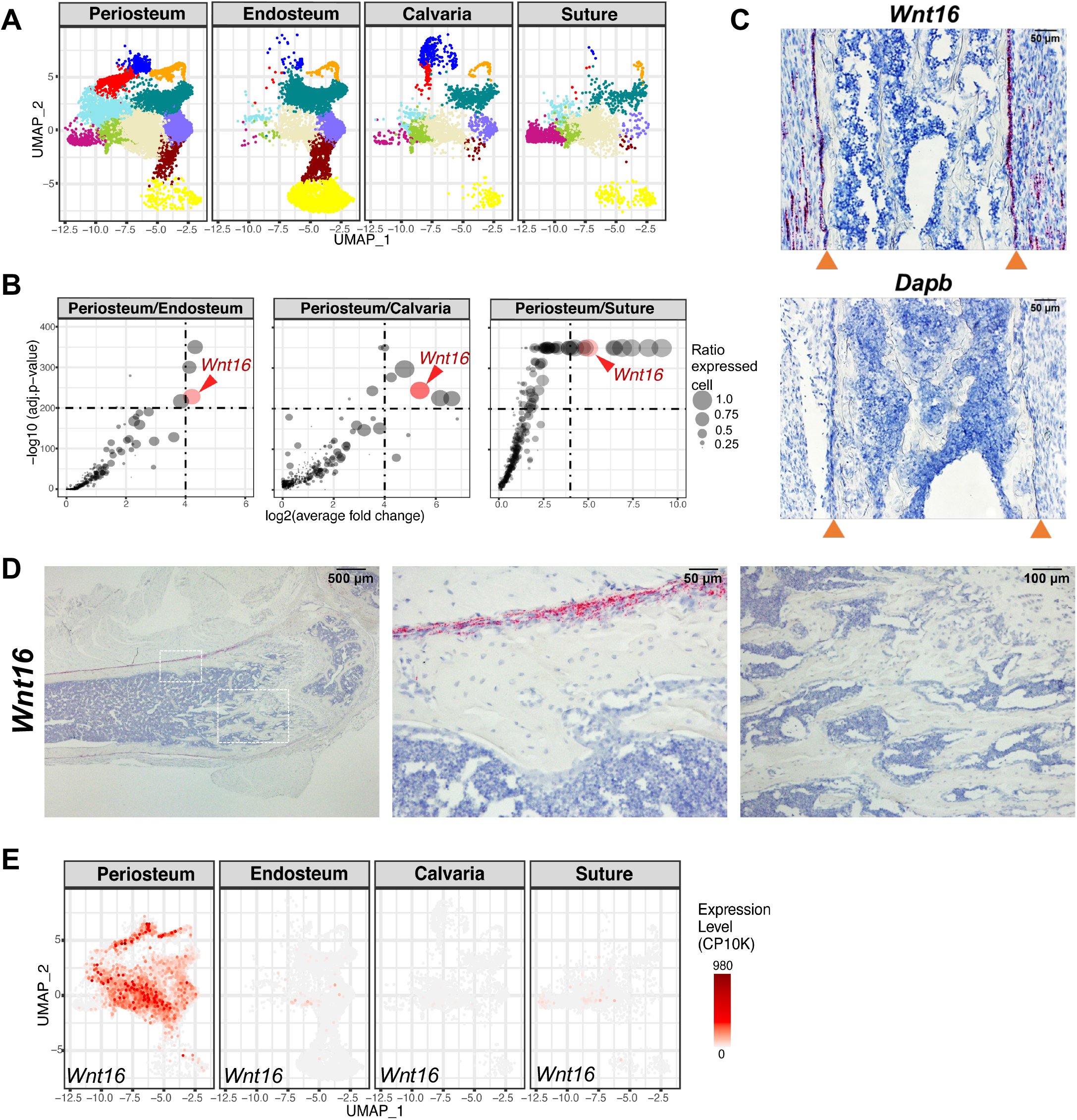
Periosteal-specific expression of *Wnt16*. **A.** UMAP representations of all the periosteal cells isolated in the present study (Periosteum), all of the endosteal and trabecular bone associated cells isolated in the present study (Endosteum), cells isolated by Ayturk et al. (24) (Calvaria), or cells isolated by Holmes et al. (25) (Suture). **B.** Differential gene expression plots comparing expression of genes in the osteo-X cluster from the indicated cell preparations. Circle size corresponds to the ratio of *Wnt16* expression between the 2 indicated bone sites. **C.** In situ hybridization using a probe for *Wnt16* and a femoral bone section from a 7-day-old male C57BL/6 mouse. Red = *Wnt16* expression. Arrowheads indicate location of periosteal bone surface. *Dapb* is a bacterial transcript used as a negative control. **D.** In situ hybridization was performed on a femoral bone section from a 4-month-old female C57BL/6 mouse. Images at the right are higher magnifications of the boxed areas in the left panel. **E.** UMAP feature plots showing expression of *Wnt16* in cells from the indicated preparations. Red = high expression.

### Relationship of osteo-X to other clusters

To better characterize cells in the osteo-X cluster, and to identify possible relationships to cells in other clusters, we sought to localize osteo-X cells in bone tissue sections using in situ hybridization with probes for transcripts highly expressed by this cluster. To facilitate orientation, we first localized osteoblasts using a *Bglap* probe, which identified osteoblasts lining much of the cancellous and endosteal bone surface, with some staining at the periosteum in 4-month-old mice (**Fig. 5A**). No staining for *Bglap* was observed in the bone marrow and little or no staining was observed in osteocytes. Two of the defining transcripts for the osteo-X cluster are *Aspn* and *Postn*. Consistent with the idea that osteo-X cells are much more abundant in periosteal tissue than at the endosteum, both transcripts were highly expressed in cells at the periosteum but in few cells within the bone marrow cavity (**Fig. 5B-C**). Strong staining was also observed in a structure with a location and morphology consistent with the groove of Ranvier (Fig. 5B-C). Neither transcript was detected in hematopoietic cells in the bone marrow (Fig. 5B-C). Notably, transcripts for *Postn* but not *Aspn* were detected in cells associated with the cancellous bone surface (Fig. 5B-C). Faint staining for both probes was detected in a subpopulation of blood vessels in the bone marrow and a few larger vessels in the diaphyseal region showed significant expression of *Aspn* (**Fig. S7**).

**Figure 5.**
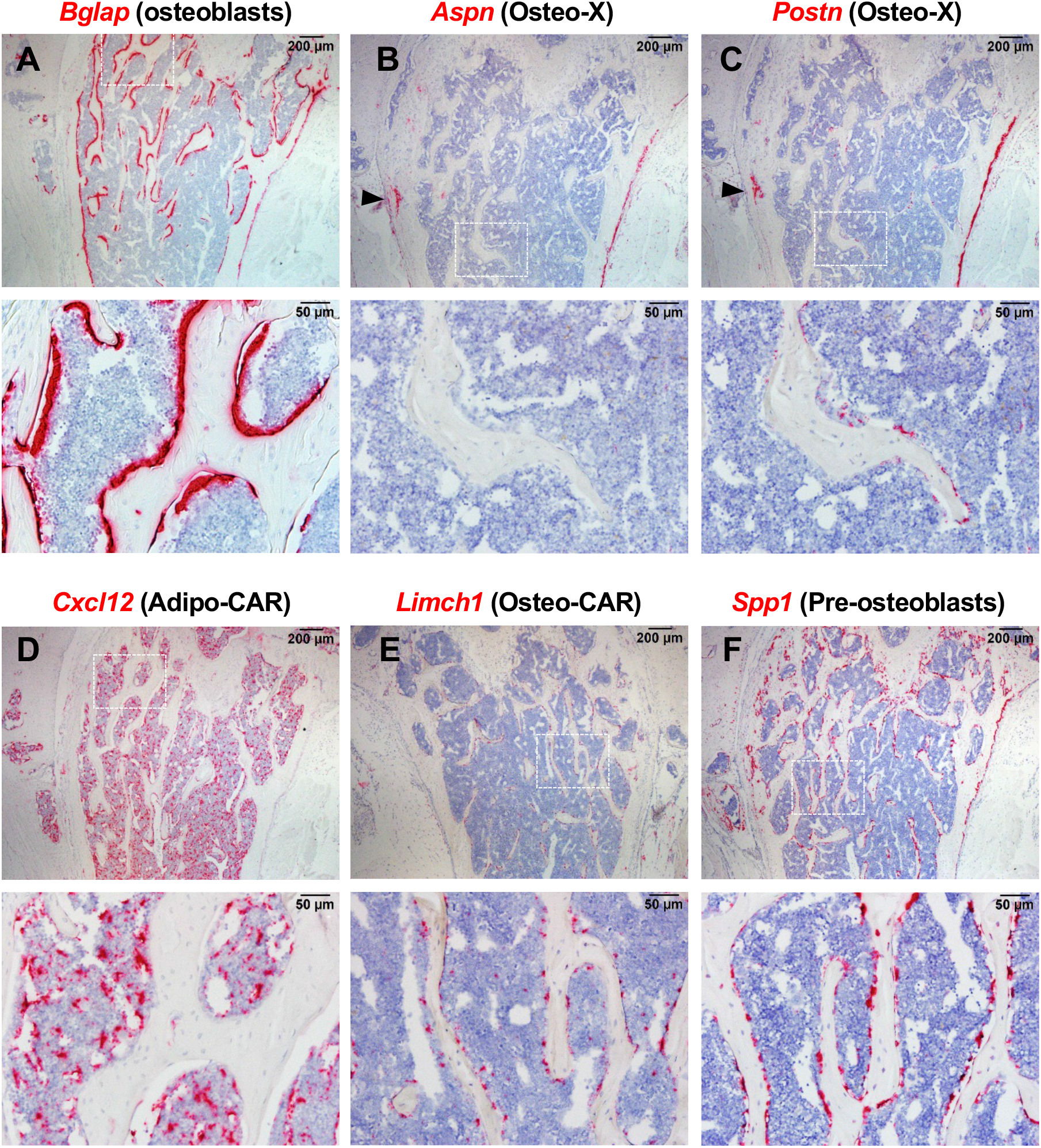
Localization of osteo-X and osteo-CAR cells. RNAScope-based in situ hybridization of femoral bone sections from a 4-month-old female C57BL/6 mouse using the probes for transcripts expressed by the following cell types: osteoblasts, defined by *Bglap* (**A**); osteo-X, defined by *Aspn* (**B**); osteo-X, defined by *Postn* (**C**), adipo-CAR, defined by *Cxcl12* (**D**); osteo-CAR, defined by *Limch1* (**E**), and pre-osteoblasts, defined by *Spp1* (**F**). Higher magnification images for each section are shown below in each panel. Red = transcript expression. Arrowheads in panel B and C indicate location of groove of Ranvier.

The presence of *Postn*-positive cells near the cancellous bone surface suggested that a subpopulation of cells in the osteo-X cluster may be in close association with either osteoblasts or their precursors. To explore this idea further, we used a probe for *Cxcl12* to detect adipo-CAR cells and a probe for *Limch1* to detect osteo-CAR cells since previous studies have shown that a population of CAR cells contributes to osteoblast formation, at least in cancellous bone (22). Consistent with earlier studies (35), *Cxcl12*-expressing cells were distributed throughout the bone marrow space, often associated with blood vessels (**Fig. 5D**). In contrast, osteo-CAR cells were much less abundant and primarily associated with the cancellous bone surface (**Fig. 5E**). Similar to osteo-CAR cells, cells expressing high levels of *Spp1*, a marker for cells we designated as pre-osteoblasts, were predominantly associated with the cancellous bone surface (**Fig. 5F**).

The close association of osteo-X (expressing *Postn*), osteo-CAR (expressing *Limch1*), pre-osteoblasts (expressing *Spp1*), and osteoblasts (expressing *Bglap*) suggests the possibility of a continuum of cells spanning from osteoblast progenitors to mature osteoblasts residing near the bone surface. If this is the case, one would anticipate that suppression of bone remodeling would eliminate mature osteoblasts but leave in place the precursors that are the target of coupling factors produced or controlled by osteoclasts (39). Therefore, to determine which, if any, of these cell populations are dependent on ongoing bone remodeling, we suppressed osteoclast formation in humanized *Tnfsf11* mice using denosumab and examined the impact on different mesenchymal cell clusters using scRNA-seq. Importantly, to ensure that our analysis of this new scRNA-seq experiment produced cluster annotations consistent with the cell clusters generated by the harmonized UMAP, we used the harmonized dataset and cluster annotation to construct a reference-based mapping model using the Azimuth pipeline (40). In essence, this approach allowed us to use the reference map to define the cell clusters in the new experiment.

We have previously shown that administration of denosumab to humanized RANKL mice once per 2 weeks for at least 4 doses potently suppresses bone remodeling (41). Therefore, we administered denosumab using this regime and then performed scRNA-seq analysis of cells associated with endosteal and cancellous bone but depleted of hematopoietic and endothelial cells to compare the mesenchymal cell distribution. We found a dramatic reduction in osteoblast number in the denosumab-treated mice (**Fig. 6A-B**), consistent with suppression of bone remodeling. The abundance of pre-osteoblasts was also slightly reduced as was expression of the pre-osteoblast marker *Spp1* (**Fig. 6B-C**). In contrast, the relative abundance of osteo-CAR and adipo-CAR cells was not dramatically altered (Fig. 6A-B). Consistent with these results, in situ hybridization revealed an almost complete loss of *Bglap*-positive osteoblasts in vertebral cancellous bone in denosumab-treated mice (**Fig. 6D**). Pre-osteoblasts, as defined by *Spp1* expression, showed robust staining on cancellous bone in the vehicle-treated mice, which was reduced but not abolished after denosumab (Fig. 6D). In contrast, the osteo-CAR marker *Limch1* and the osteo-X marker *Postn*, displayed little change after denosumab administration (Fig. 6D). Together, these results demonstrate that osteoblasts, and to some extent pre-osteoblasts, are highly dependent on bone resorption and coupling factors for their existence in cancellous bone. However, the abundance of osteo-CAR and osteo-X cells, and their close association with the cancellous bone surface, do not require on-going bone remodeling.

**Figure 6.**
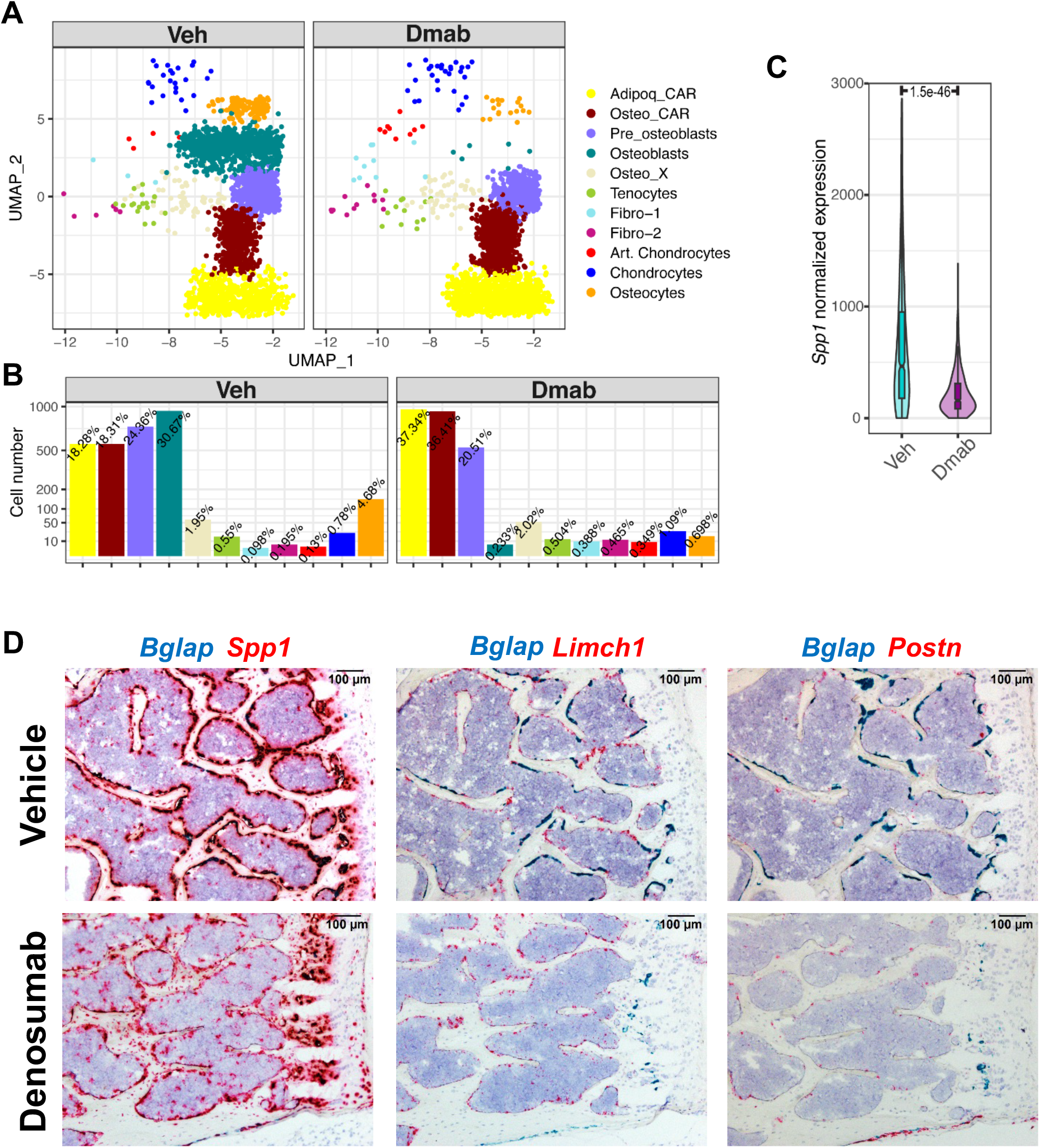
Suppression of remodeling does not alter location of osteo-X and osteo-CAR cells. **A.** UMAP representations of mesenchymal cells isolated from endosteal and trabecular bone by negative selection from mice treated with vehicle (Veh) or denosumab (Dmab). Cell names and color codes are the same as in earlier figures. **B.** Bar graphs indicating the relative abundance of the cell clusters shown in panel A. **C.** Violin plot of Spp1 expression in cells of the pre-osteoblast clusters shown in panel A. **D.** Two-color RNAScope in situ hybridization using the indicated probes and vertebral bone sections from mice treated with vehicle or denosumab.

While osteo-X and osteo-CAR exhibit characteristics consistent with a role as osteoblast progenitors, they do not possess commonly accepted characteristics of mesenchymal stem cells or skeletal stem cells, such as expression of *Ly6a* (Sca1) and *Cd34* (Fig. S4). Cells in the cluster that we have designated fibro-2 do express these genes and have been proposed by others to represent early mesenchymal progenitors analogous to PαS cells (21). To determine the location of such cells in adult remodeling bone, we performed in situ hybridization with probes for *Clec3b* and *Mfap5*, two transcripts highly expressed in fibro-2 cells along with *Ly6a* and *Cd34*. We selected *Clec3b* and *Mfap5* rather than *Ly6a* and *Cd34* since the latter two genes are also highly expressed by endothelial progenitor cells (42). Both transcripts were easily detected in cells in the outer layers of the periosteum, consistent with the presence of early mesenchymal progenitors at this site observed by others (**Fig. S8**) (43). In contrast, we did not detect cells expressing either transcript within the bone marrow cavity or associated with cancellous bone or blood vessels (Fig. S8). These results suggest that the cells represented by the fibro-2 cluster reside predominantly, if not exclusively, at the periosteum.

## Discussion

With the advent of single cell RNA sequencing, the characteristics that define a cell type, or the number of different cell types, associated with a particular tissue are becoming increasingly difficult to agree upon (44). One benefit of our harmonization approach is that it allowed us to identify what may be a minimum number of general mesenchymal cell types associated with murine bone. An important caveat to this view is that our approach can only identify cell types that can be sequenced using the 10X Genomics platform. For example, cells with a profile consistent with mature bone marrow adipocytes were not present in any of the datasets that we used and this is likely due to an inability of such cells to survive the isolation and sequencing methods. Even with this limitation, our approach may facilitate future sequencing projects by providing marker genes that can be used to guide consistent cluster identification.

The heterogeneity within many of the clusters identified here indicates that they contain multiple different cell types. The chondrocyte cluster, for example, clearly contains resting, proliferating, and hypertrophic, or perhaps pre-hypertrophic, growth plate chondrocytes. Similarly, close examination of the heatmaps for each cluster reveals multiple distinct patterns of gene expression within each cluster. Whether most of these differences represent distinct differentiation states or whether they represent a given cell type in different modes of function will require significant additional study and agreement within the field.

The significance of the restriction of *Wnt16* expression to the periosteum is unclear. Periosteal-specific expression of *Wnt16* is consistent with its demonstrated role in periosteal but not cancellous bone formation (45, 46). It is noteworthy that many of the cell types expressing *Wnt16* at the periosteum are also present within the bone marrow cavity. Expression specifically at the periosteum suggests that a signal stimulating *Wnt16* production exists exclusively in the periosteal environment. Accumulating evidence suggests that skeletal muscle directly promotes bone formation (47, 48) and it is possible that muscle cells provide a signal that stimulates *Wnt16* expression in the periosteal environment. Regardless of the mechanism underlying the restricted expression of *Wnt16*, its production specifically in periosteal cells provides a useful genetic marker for the presence of periosteal cells within scRNA-seq datasets. We were unable to detect *Wnt16*-expressing cells by in situ hybridization at the endosteum, on cancellous bone, or within the bone marrow cavity in either growing or adult mice. Thus, any bone-associated cells expressing this gene in scRNA-seq datasets are likely derived from periosteal tissue.

The identity and location of skeletal stem cells or mesenchymal stem cells remain active areas of debate and investigation. In our analysis, the only mesenchymal cell cluster to express accepted markers of mesenchymal stem cells, such as *Ly6a* (Sca1) and *Cd34*, is fibro-2. Fibro-2 cells, as defined by expression of *Clec3b* and *Mfap5*, were easily detected at the periosteum and surrounding tissue but not within the bone marrow cavity. This finding suggests that osteoblast progenitors within the bone marrow cavity of adult mice may not be derived from cells with classical stem cell markers. This scenario is consistent with previous reports suggesting that the majority of osteoblasts produced during physiological bone remodeling are derived from committed replicating progenitors rather than stem cells (49, 50).

What is the relationship, if any, of either adipo-CAR or osteo-CAR cells to osteoblasts? Lineage tracing studies in adult mice suggest that at least a portion of osteoblasts on cancellous bone are derived from some type of CAR cell. Specifically, transient activation of a Cxcl12-CreERT2 transgene at 2 months of age led to labeling of more than 30% of trabecular osteoblasts and this labeling remained constant until at least 18 months of age (22). Notably, cortical osteoblasts were not labeled at any time in these same mice. Similarly, Adipoq-Cre targets almost all CAR cells, as defined by Cxcl12-GFP transgene expression, and while few osteoblasts were labeled at 4 weeks of age, about 20% were labeled at 6 months of age (51). Whether the osteoblasts labeled in these studies were derived from adipo-CAR cells, osteo-CAR cells, or both is unclear. Nonetheless, the long-term labeling of osteoblasts by transient activation of the Cxcl12-CreERT2 transgene suggests that some of the CAR cells that it targets represent self-renewing osteoblast progenitors in adult remodeling bone.

In the present study, we localized osteo-CAR cells, as defined by *Limch1* expression, to the endosteal and trabecular bone surface of adult mice. Further, we showed that in actively remodeling bone, these cells are associated with osteoblasts on the bone surface and that they overlap with or are associated with pre-osteoblasts, as defined by *Spp1* expression. Strikingly, suppression of bone remodeling using denosumab led to the elimination of osteoblasts and suppression of *Spp1* expression in pre-osteoblasts, but no change in the abundance of osteo-CAR cells, which remained in close association with the bone surface. Based on their location, their independence from bone remodeling, and the lineage-tracing studies noted earlier, we suggest that osteo-CAR cells, or a subpopulation of osteo-CAR cells, represent a continuous source of osteoblast progenitors in adult remodeling bone.

The study that initially proposed the term “osteo-CAR” to refer to cells expressing both *Cxcl12* and osteoblast-lineage genes reported that they were localized to arteriolar and non-vascular niches (18). In contrast, our study localized these cells primarily to regions near endosteal and trabecular bone surfaces. As noted by the authors of this earlier work, the extreme signal intensity of the alkaline phosphatase staining that they used to identify osteoblastic cells may have prevented detection of CAR cells near the bone surface (18).

Our rationale for proposing that osteo-CAR, rather than adipo-CAR cells, serve as osteoblast progenitors is based largely on the close proximity of osteo-CAR cells to the bone surface and to existing teams of osteoblasts. Transient activation of an Osx1-CreERT2 transgene at E14.5 labeled a population of osteoblast progenitors that remained spatially-restricted to the region of the femur that existed at the time of labeling even up to 10 months of age (52). This later finding reveals that, during normal bone remodeling, osteoblasts are derived from progenitors that do not migrate large distances, suggesting that osteo-CAR cells are better positioned to serve as osteoblast progenitors than the majority of adipo-CAR cells.

If Osteo-CAR cells do represent an important source of osteoblasts in remodeling bone, it might be expected that deletion of beta-catenin using Dmp1-Cre would inhibit osteoblast formation due to the fact that Dmp1-Cre targets CAR cells. However, it is important to note that Dmp1-Cre targets only about 30% of CAR cells, as defined by Cxcl12-GFP transgene expression (53). Therefore, loss of beta-catenin in only 30% of osteoblast progenitors would not be expected to completely inhibit osteoblast formation, which is consistent with the phenotype of mice lacking beta-catenin in Dmp1-Cre targeted cells (54).

As opposed to life-long production of osteoblasts by a single type of resident mesenchymal or skeletal stem cell, evidence is accumulating that osteoblasts are derived from a variety of different progenitors depending on the age of the animal, the location of the osteoblasts in the skeleton, and the need for repair (28). Results presented herein suggest that osteo-X and osteo-CAR cells represent potential sources of osteoblast progenitors located in close proximity to the endosteal and cancellous bone surfaces in adult mice. The developmental origin of osteo-X and osteo-CAR cells will require additional study but a likely source is the cell type that is marked by an Osx1-CreERT2 transgene and that gains access to the bone interior at an early stage of endochondral bone formation (55). While initially associated with invading blood vessels, such cells may give rise, over time, to progenitors residing near the bone surface. The presence of osteoblast progenitors near the bone surface is consistent with the ability of osteocyte-derived sclerostin to inhibit osteoblast formation as sclerostin function depends on local membrane retention by Lrp4 (56). It will be important in future studies to determine whether osteo-X and osteo-CAR cells do in fact serve as osteoblast progenitors and whether their ability to do so is altered in conditions that lead to low bone formation.

## Acknowledgments

We thank the staff of the UAMS Department of Laboratory Animal Medicine for technical support. This work was supported by the National Institutes of Health (R01AR049794 and P20GM125503), the Veterans Administration (I01 BX000294), and by UAMS tobacco settlement funds.

## Author Contributions

I.N., J.X., M.A., and C.A.O. conceived and designed the studies. I.N., J.X., M.O., C.B-G, V.W., Q.F., H-N.K., and C.A.O. performed experiments and analyzed data. C.A.O. wrote the manuscript. All authors edited and approved the final version of the manuscript.

## Declaration of Interests

The authors declare no competing interests.

## Methods

### Animal housing and care

Dmp1-Cre, Osx1-Cre, Ai9, and humanized Tnfsf11 mice have been described previously (12, 27, 57, 58). Wild type C57BL/6J mice were obtained from the Jackson Laboratory. Mice were socially housed at 2-5 animals per cage using a blend of 1/4″ corncob bedding and white enrichment paper both produced by Andersons Incorporated. The animal colony was specific pathogen free based on the Division of Laboratory Animal Medicine’s exclusion list. Mice were provided ad libitum water and an irradiated Purina diet of either 5V5M (for breeders) or 5V5R (for maintenance). The temperature range in the room was 68-79 F with a set point of 71 +/- 2, additionally the humidity range was 30%-70%. The room was on a 12:12 hour light cycle and the illumination was 364 lux measured 1 meter from the floor.

### Study approval

All animal procedures were reviewed and approved by Institutional Animal Care and Use Committee of the University of Arkansas for Medical Sciences.

### Cell isolation

For all experiments using mice harboring the Cre driver transgenes and the tdTomato reporter allele (Ai9), mice between the ages of 8 and 14 weeks were used. To obtain cells associated with endosteal and trabecular bone, femurs and tibias from a single mouse were cleaned of soft tissues and the periosteum removed by scraping with a scalpel. After removing the epiphyses, bone shafts were cut open lengthwise using a scalpel and bone marrow cells were removed by flushing with PBS containing 1% BSA. The remaining bone fragments were then cut into smaller pieces (approximately 1 mm in length) and placed into a single well of a 6-well culture dish containing Hank’s Balanced Salt Solution (HBSS) and 2 Wunsch units of LiberaseTM (MilliporeSigma). The cells were incubated for 20 minutes at 37C with shaking after which the released cells were collected by pipetting and incubated on ice. The bone fragments were digested a second time for 20 minutes with LiberaseTM and the cells saved as above. The bone fragments were then incubated in PBS containing 5 mM EDTA and 0.1% BSA for 20 minutes at 37C with shaking. The released cells were collected and saved as above. The bone fragments were subsequently exposed to alternating exposures of either LiberaseTM or EDTA to yield a total of 4 LiberaseTM digestions and 3 EDTA incubations. All fractions were pelleted at 300xg for 10 minutes and resuspended in 0.5 ml PBS containing 1% BSA. Fractions were then pooled, filtered through a 70 µm filter, pelleted at 300xg for 10 minutes, and resuspended in less than 1 ml sorting buffer (PBS containing 0.5% BSA and 2 mM EDTA).

To obtain cells from the periosteum, intact femurs were dissected from 2 mice. Muscles were carefully removed using a scissors without damaging the periosteum. The femurs were placed into a 50 ml conical centrifuge tube containing HBSS with calcium and 2 Wunsch units of LiberaseTM (MilliporeSigma) and incubated for 20 minutes at 37°C with shaking. The released cells were collected by pipetting and stored in PBS containing 1% BSA on ice. Bones were then washed twice with 50 ml of PBS and incubated in calcium free PBS with 5 mM EDTA and 0.1% BSA and incubated for 20 minutes at 37°C with shaking. The released cells were collected and stored on ice. The femurs were digested one more time for 20 minutes with LiberaseTM and the cells were collected. All cells collected were then pooled, filtered through a 70 µm filter, and pelleted at 300xg for 10 minutes. Cells were then resuspended in 200 µl sorting buffer for sorting.

Cells from mice harboring the Ai9 allele were sorted using a BD FACS Aria III with a 100 µm nozzle to collect tdTomato-positive cells. For mice not harboring a reporter allele, cells were obtained by LiberaseTM and EDTA incubation as above but instead of isolation by FACS, cells were depleted of hematopoietic cells and endothelial cells using a lineage depletion kit (Miltenyi Biotec, cat. no. 130-090-858) followed by CD45, CD117, and CD31 microbeads (Miltenyi Biotec, cat. nos. 130-0520301, 130-091-224, and 130-097-418) according to the manufacturer’s instructions. Cells in the flow-thru were pelleted at 300xg for 10 minutes and resuspended in 20 µl of sorting buffer.

### 10X Genomics sequencing

Single cells isolated from digested bones were stained with ReadyProbes™ Cell Viability Imaging Kit, Blue/Green (Thermo Fisher Scientific, catalog # R37609), and manually counted using hemocytometer under EVOS M7000 microscope (Thermo Fisher Scientific). Immediately following cell counting, samples were processed using Chromium Next GEM Single Cell 3’ Reagent Kits v3.1 (Dual Index) as described in the manufacturer’s instructions (10X Genomics). In brief, aiming for 8000 cells per library, single cell suspensions with more than 70% live cells were loaded onto Chromium Controller (10X Genomics) to generate gel beads-in-emulsions. Then co-partitioned cells were lysed, primers were released from Gel Beads, and barcoded full-length cDNA was produced and amplified. 3’ Gene expression libraries were generated from cDNA by fragmentation, end repair, A-tailing, adaptor ligation and index PCR amplification. The concentration and size distribution of final libraries were assessed by Qubit™ 1X dsDNA HS Assay (Thermo Fisher Scientific, catalog # Q33231) and the Fragment Analyzer System (Agilent). Libraries were sequenced either on a NextSeq500 or NovaSeq 6000 (Illumina) with paired-end mode (read1: 28 cycles, read 2: 90 cycles, i7: 10 cycles, i5: 10 cycles) to generate a minimum of 20,000 read pairs per cell.

### Bioinformatic analysis of single cell RNA-sequencing

The fastq files were preprocessed using Cell Ranger software version 6 (10X Genomics) to produce feature-barcode matrixes. The alignments were performed using mouse reference genome mm10. The feature-barcode matrixes were imported for further analysis in R suite software using Surat package version 4.2.0 (40). Cells with between 500 and 3000 transcripts were included for further analysis. The harmonization between samples was performed using a canonical correspondence analysis method based on the top 50 principal components and 6000 most variable features to minimize batch effect. The harmonized results were used for clustering using Louvain algorithm with multilevel refinement and Uniform Manifold Approximation and Projection (UMAP) for dimension reduction. The gene specific markers of individual clusters we and differential expressed genes identified using MAST algorithm for cell type identification (59). The normalized expression kernel-weighted density values were calculated for selected marker genes using Nebulosa (60) then visualized on either UMAPs or violin-boxplots.The expression level of the selected genes was plotted based on normalized expression value of relative count per 10,000 (CP10K). The integrated datasets were used as the input to Azimuth workflow (40) to construct a reference map and the annotated cell type models for reference-based mapping analysis of denosumab administration experiments.

### Denosumab administration

Denosumab (Prolia®, 60 mg/ml) was purchased from the UAMS pharmacy, aliquoted into 0.5 ml sterile microtubes, and stored at 4°C protected from direct light. Mice were injected subcutaneously with either vehicle (saline) or denosumab at a dose of 10 mg/kg body weight, in a total volume of 100 µl, once every two weeks (Q2W) for either 4 or 5 doses. In both cases, bones were harvested 2 weeks after the final dose.

### RNAscope

Murine femurs and lumbar spines were processed for paraffin sectioning by first fixing in Millonig’s 10% buffered formalin for 40 hr at 4C. Bones were then decalcified in 14% EDTA for one week, after which they were dehydrated into 100% ethanol and embedded in paraffin for sectioning. All aqueous solutions were prepared with DEPC-treated water. RNA in situ hybridization was performed using the RNAscope 2.5 HD detection reagent RED (cat. no. 322360) and Duplex (cat. no. 322500) kits, following the manufacturer’s instructions (Advanced Cell Diagnostics, Newark, CA). In brief, eight-micron paraffin sections were incubated at 60°C for 60 minutes, deparaffinized, and pretreated with RNAscope Hydrogen Peroxide (cat. no. 322335) for 10 minutes at RT. Sections were then incubated in the custom pretreatment solution (cat. no. 300040) at 40C for 15-45 minutes. The following probes were incubated on the sections for 2h at 40°C and detected with RNAscope 2.5 AMP 1-6 for RED kit and AMP1-10 for Duplex kit: Dapb (cat. no. 310043), Bglap (Mm-Bglap cat. no. 478941), Limch1 (Mm-Limch1 cat. no. 591801), Spp1 (Mm-Spp1 cat. no. 435191), Clec3b (Mm-Clec3b cat. no. 539561), Mfap5 (Mm-Acp5 cat. no. 465001), Postn (Mm-Postn cat. no. 418581), Aspn (Mm-Aspn cat. no. 502051), Wnt16 (Mm-Wnt16 cat. no. 401081), Cxcl12 (Mm-Cxcl12 cat. no. 422711), and Cdh5 (Mm-Cdh5 cat. no. 312531). Hybridization time for AMP 5, 6, 9, and 10 were modified for each probe. Signal was detected for 10 min at RT. Sections were counterstained with hematoxylin, dehydrated at 60°C for 20 minutes, and then mounted in VectaMount permanent mounting medium (Vector Laboratories, Newark, CA).

## Supplementary Figures

**Figure S1. Initial cluster analysis of Dmp1-Cre and Osx1-Cre targeted cells. A.** Diagram of the approach used to perform single-cell RNA-seq of cells from Dmp1-Cre;Ai9 and Osx1-Cre;Ai9 mice in this study. **B.** Initial UMAP representations of cell clusters (0–18) targeted by Dmp1-Cre or Osx1-Cre in endosteal and trabecular bone (Endo) or in periosteal bone (Peri). **C.** The left panel is a combined UMAP of all the cells in panel B with color-coded clusters (0–18) and the UMAP feature plots on the right show expression of the indicated transcripts. Red = high expression. **D.** UMAP feature plots showing levels of the tandem-dimer tomato mRNA, produced by the activated reporter gene in Ai9 mice, in each of the cell preparations shown in panel A. Red = high expression.

**Figure S2. Cluster analysis of combined datasets.** The top panel is a UMAP representation all cell clusters identified in the combined dataset consisting of the cells isolated from Dmp1-Cre and Osx1-Cre mice in this study as well as cells isolated in 9 published studies. The bottom panels are UMAP feature plots showing expression of the indicated genes that are characteristic of specific cell types: *Ptprc* (encoding Cd45) – hematopoietic cells; *Pdgfra* – mesenchymal cells; *Cdh5* – endothelial cells; *Myh11* – pericytes; *Mog* – Schwann cells; *Cd79b* – B lymphocytes; *Cd3g* – T lymphocytes; *S100a9* – myeloid cells. Red = high expression.

**Figure S3. Comparison of cell yields for different studies.** Bar graphs showing the relative and absolute abundance of different cell types isolated in datasets from published studies used in here as well as the cells isolated from Dmp1-Cre and Osx1-Cre mice in the current study. (endo) = efforts were made to remove periosteal cells from long bones before digestion with collagenase.

**Figure S4.** UMAP feature plots of the indicated transcripts in mesenchymal cells of the combined dataset. Red = high expression.

**Figure S5.** UAMP feature plots of the indicated genes in mesenchymal cells of the combined dataset. Red = high expression.

**Figure S6. Polar plot of osteocyte marker genes.** The top osteocyte marker genes identified from our study were compared with previously reported osteocyte marker genes by others Agoro et al.(37), Youlten et al.(38), and Wang et al (26). Only genes that were identified in at least two of the four studies were included in the polar plot.

**Figure S7. Localization of osteo-X cells and blood vessel endothelial cells.** Two-color RNAScope-based in situ hybridization using femoral bone sections from a 4-month-old female C57BL/6 mouse. Probes for *Aspn* and *Postn* (red) are representative of osteo-X cells and the probe for *Cdh5* (teal) is representative of blood vessel endothelial cells. Higher magnification images of metaphyseal regions of the top panels are shown below them.

**Figure S8. The bone marrow cavity contains few cells with mesenchymal stem cell markers.** Single color RNAScope-based in situ hybridization of femoral bone sections from a 4-month-old female C57BL/6 mouse. The identity of the probes is shown at the left. Red = transcript expression.

## Notes

### Competing Interest Statement

The authors have declared no competing interest.

