## Supplementary figures and images for "A framework for defining mesenchymal cell types associated with murine periosteal and endosteal bone"

### Suplementary

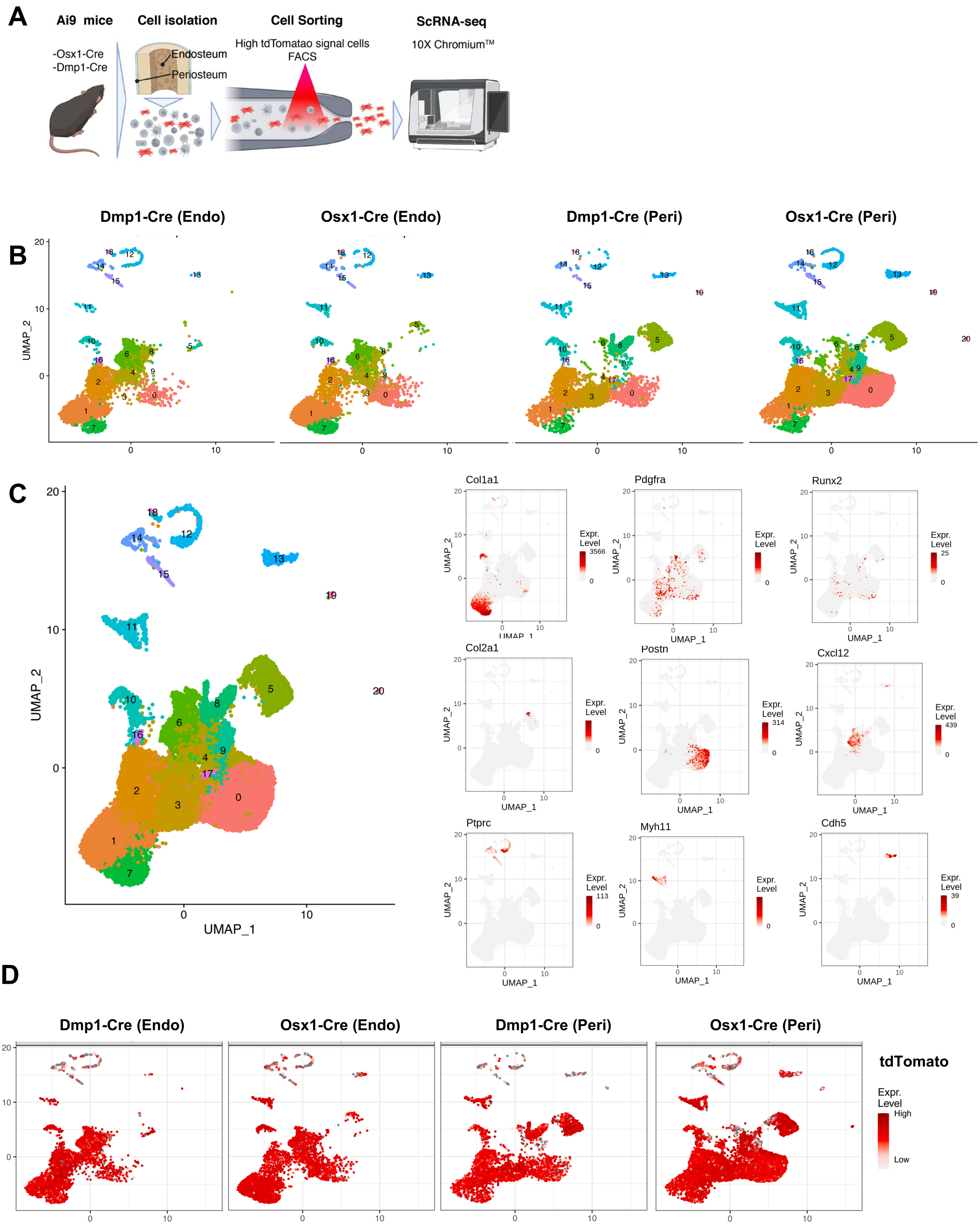

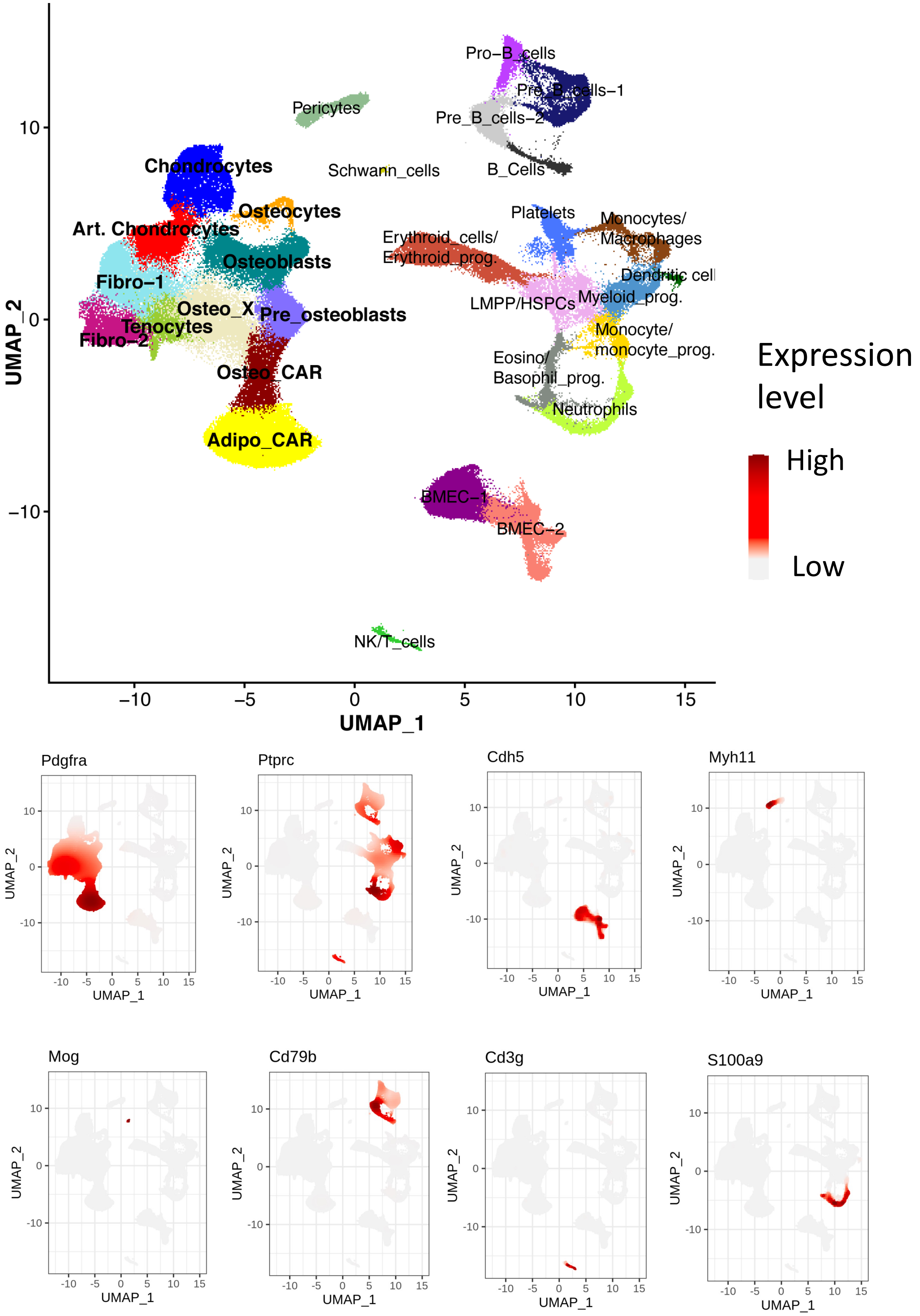

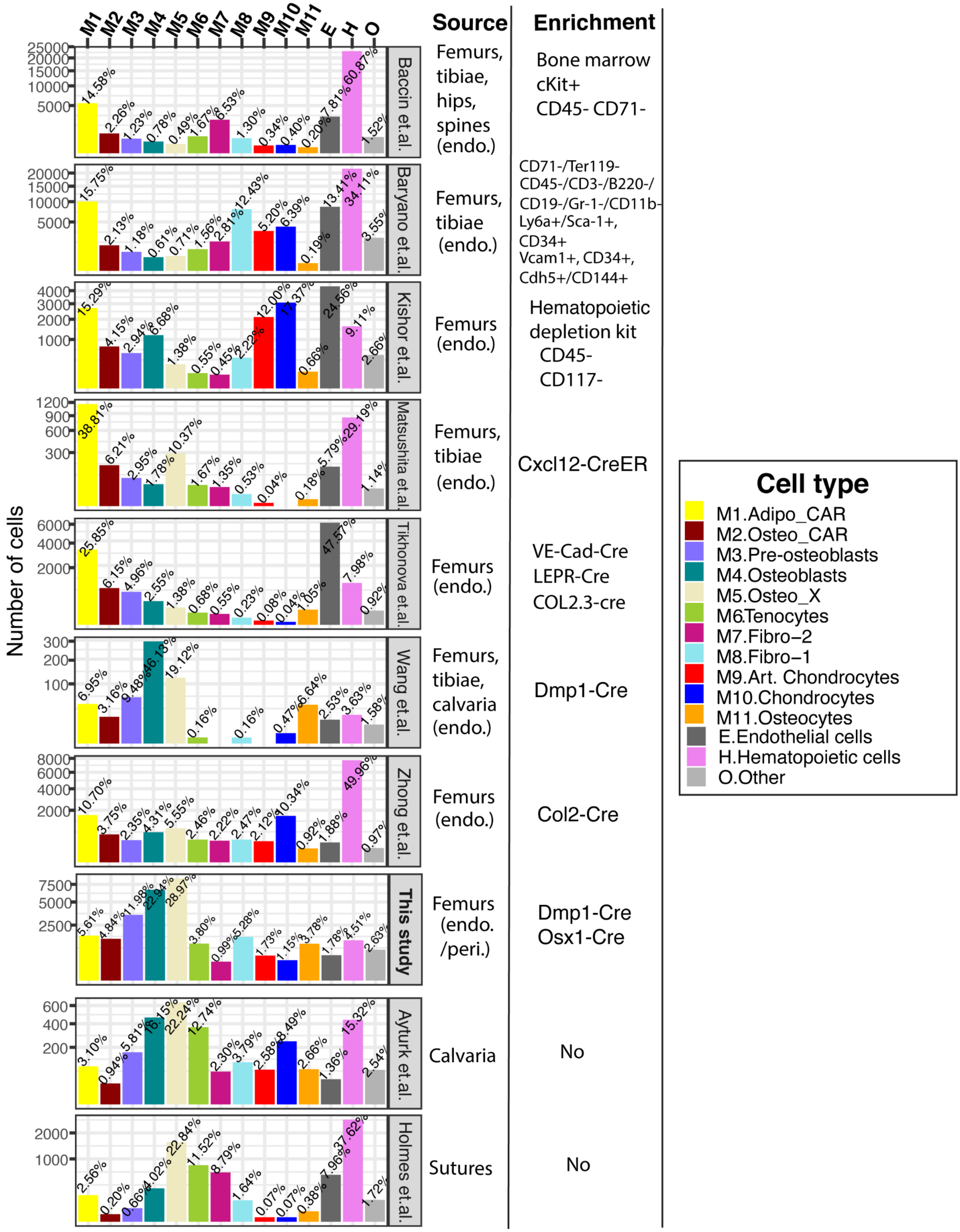

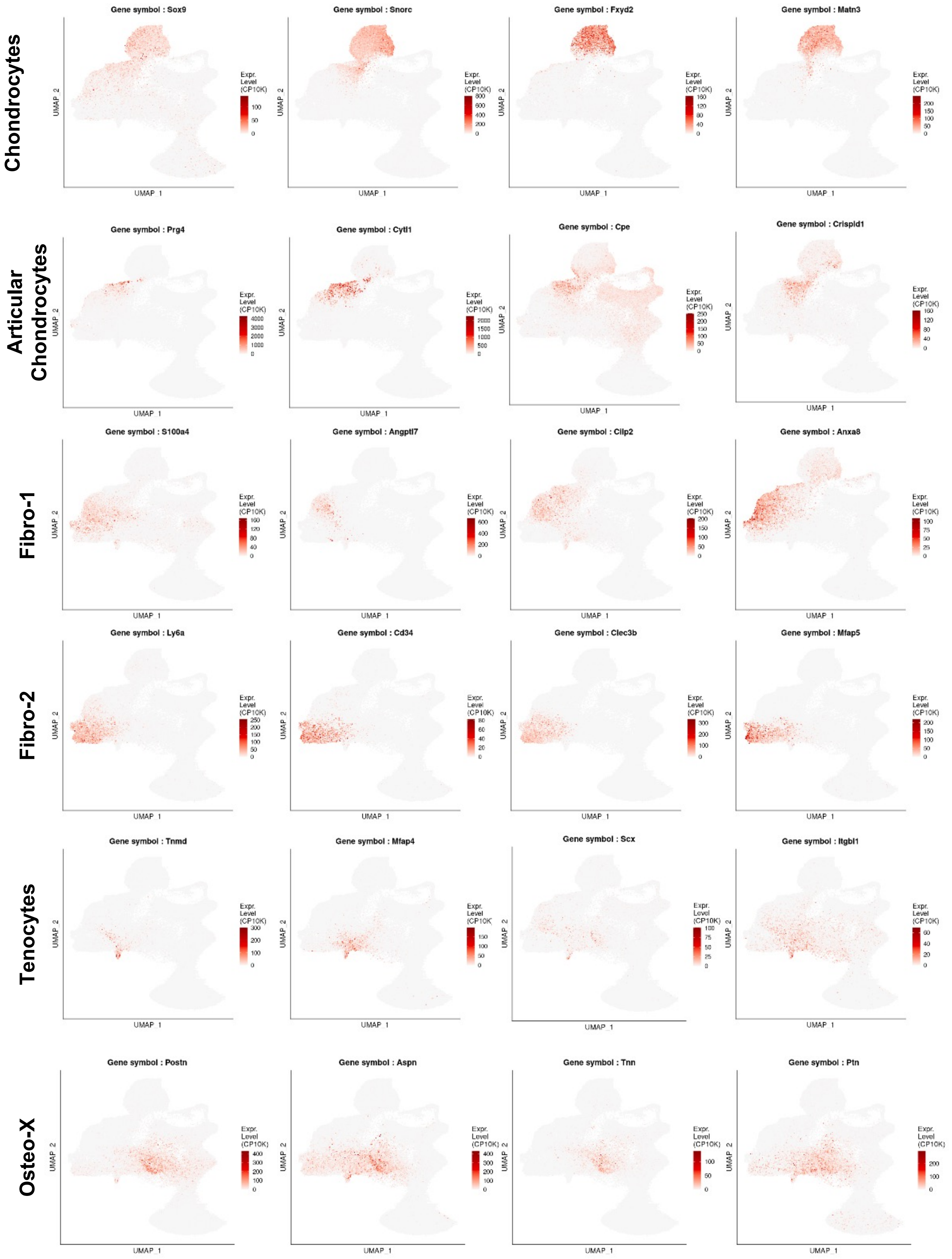

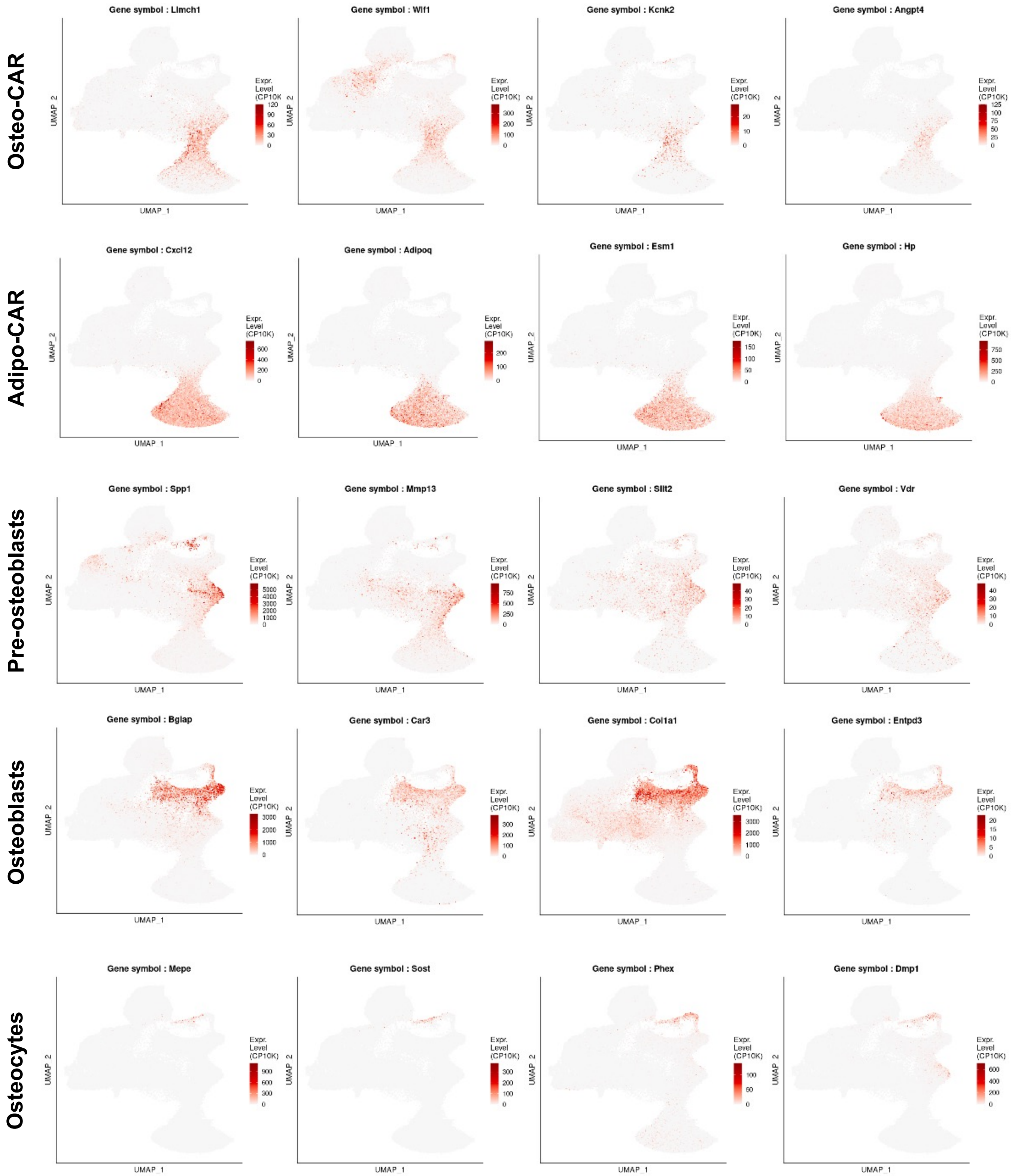

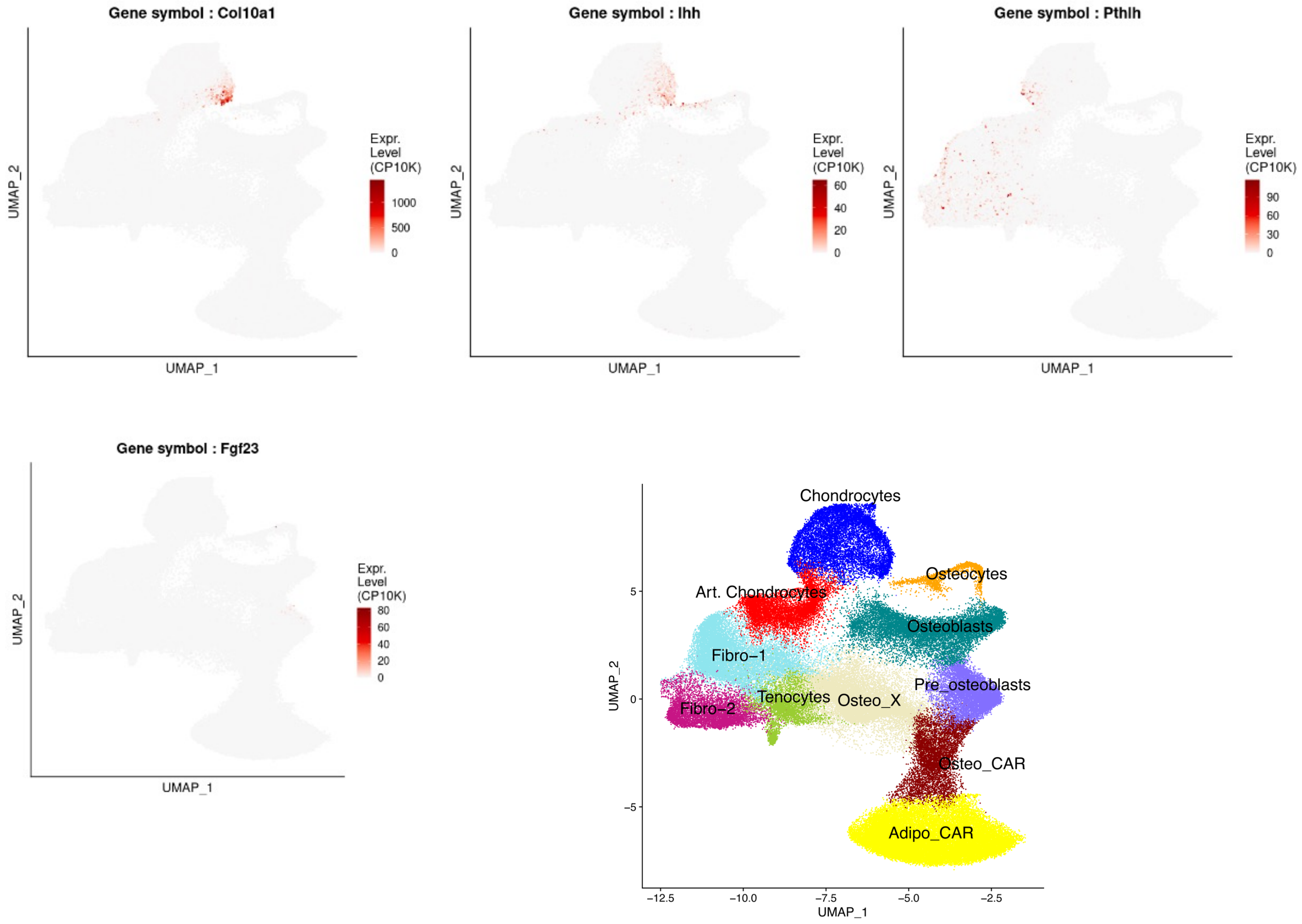

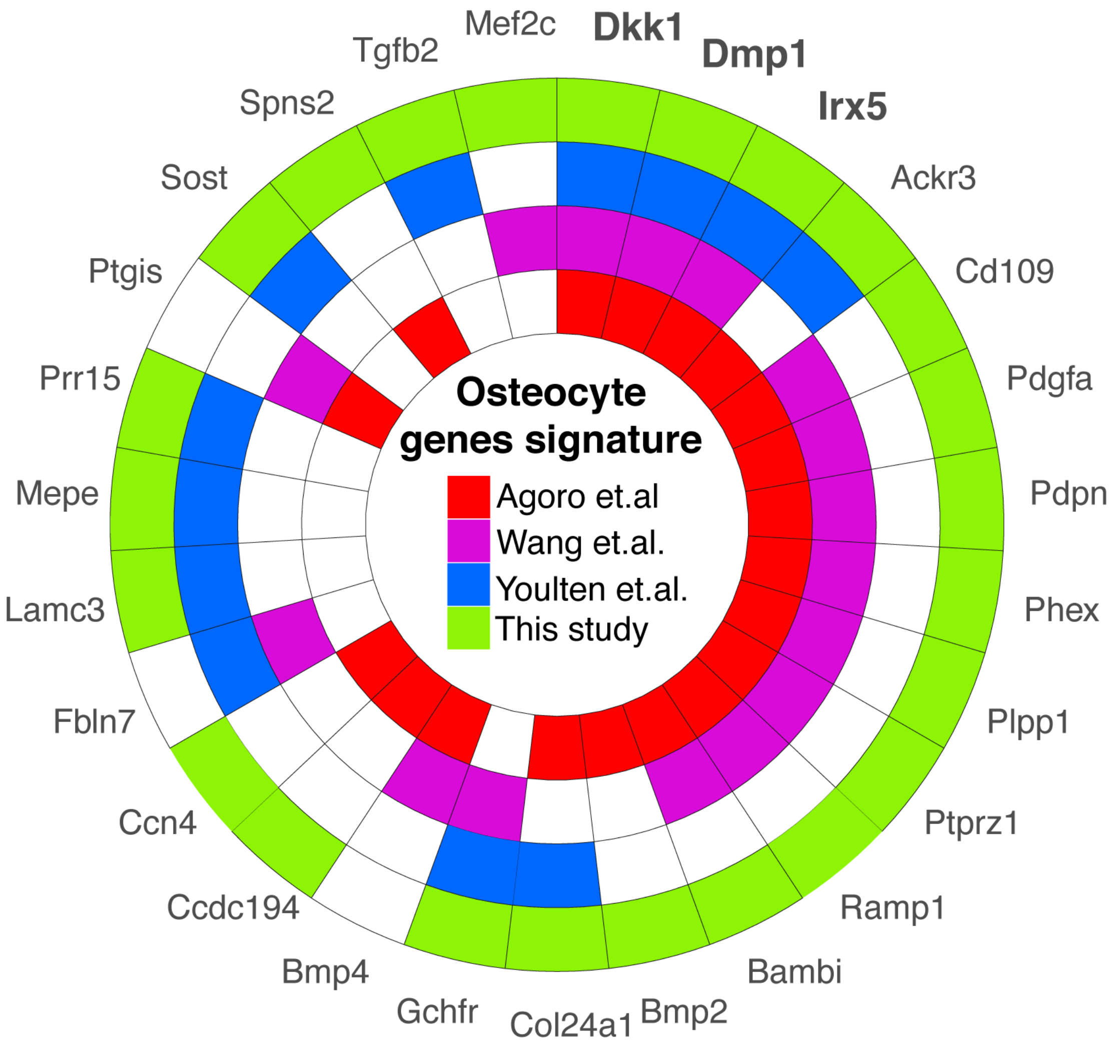

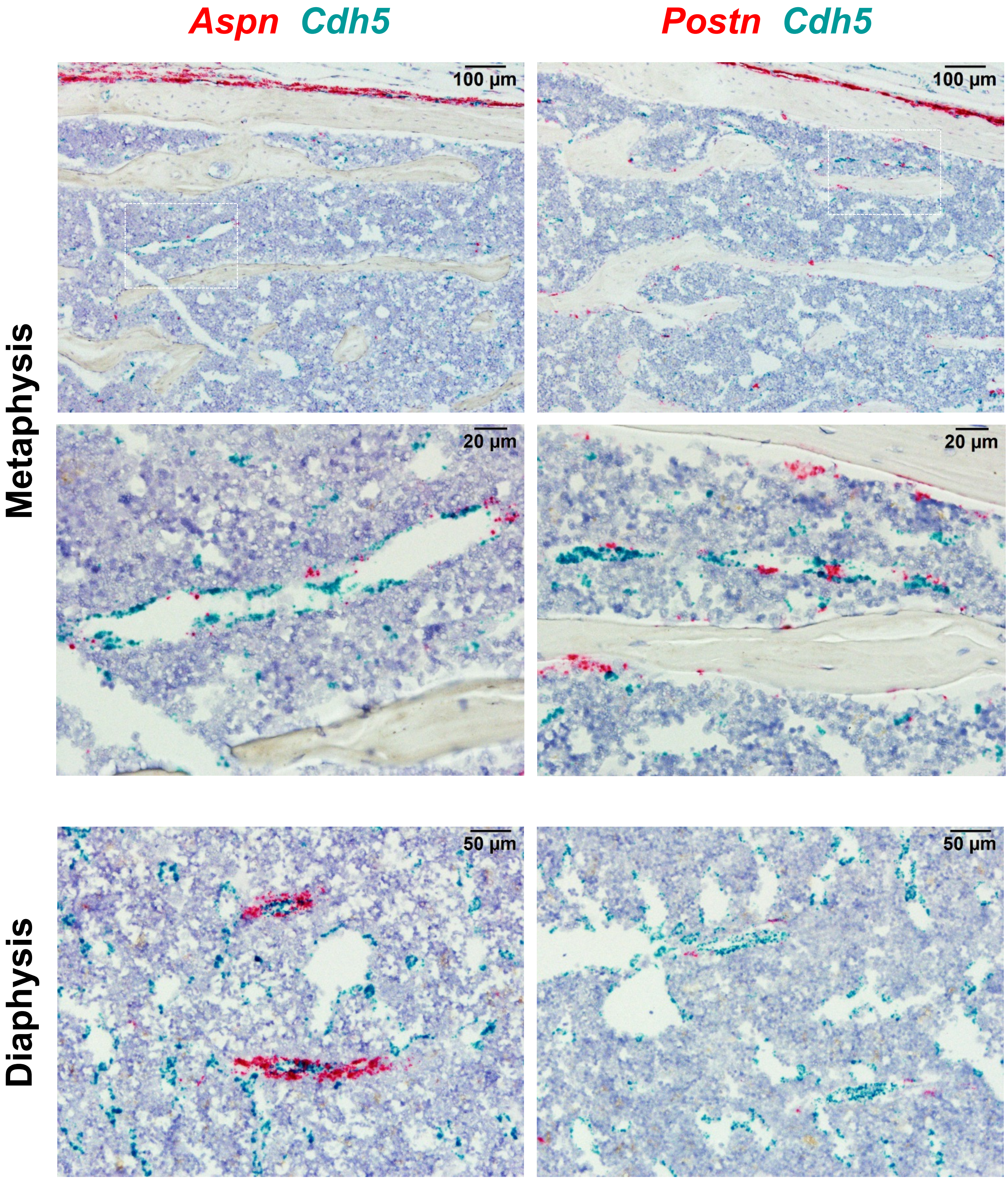

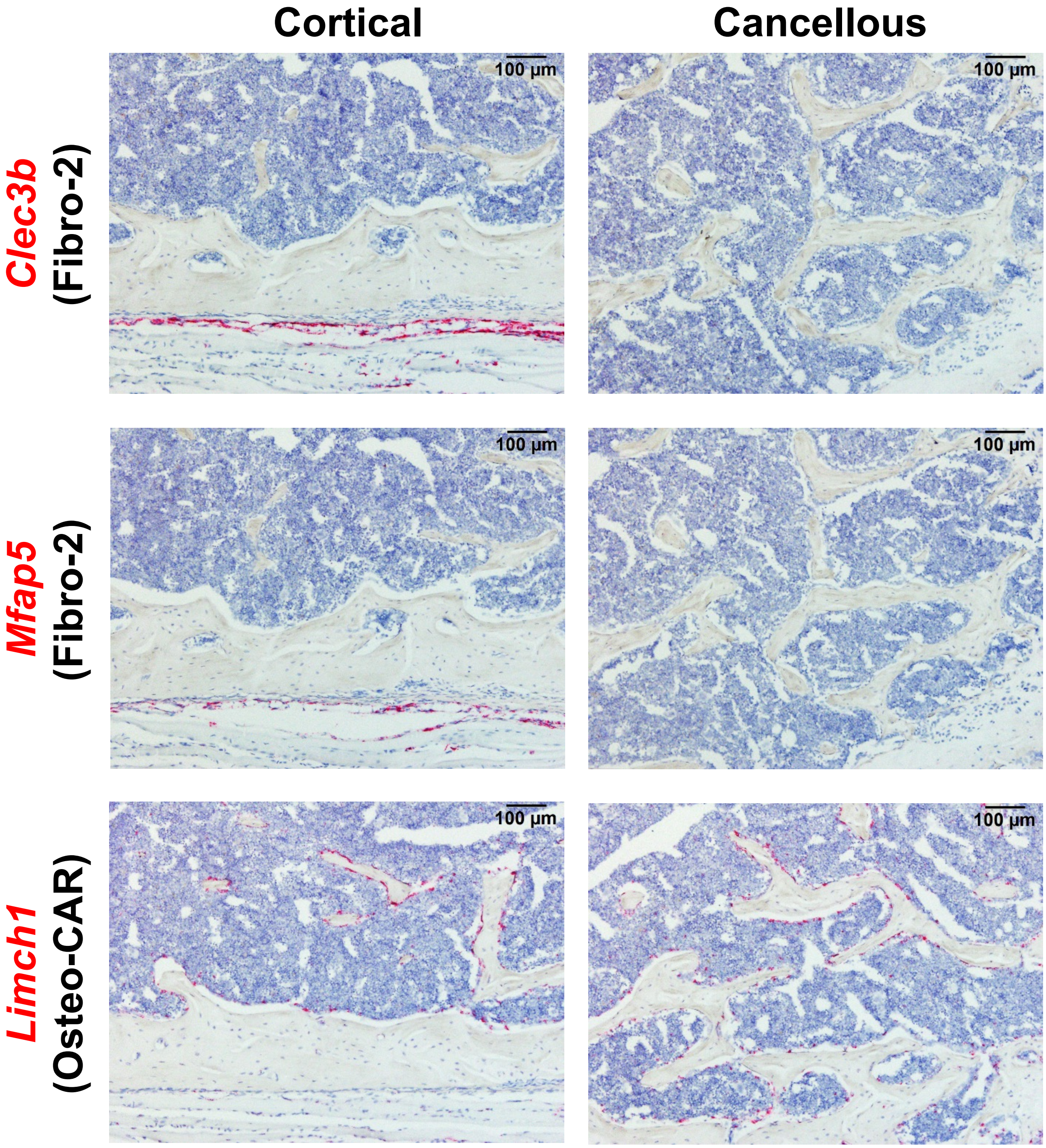
